# Development of antibody-drug conjugates targeting L1CAM to treat metastatic cancer

**DOI:** 10.1101/2025.10.01.679843

**Authors:** Jin Suk Park, Carson Kenum, Lan He, Abdul G. Khan, Mary Ann Pohl, Thomas E. White, Sreekumar R. Kodangattil, Charles M. Rudin, Paul J. Balderes, Ivo C. Lorenz, Joan Massagué, Karuna Ganesh

## Abstract

Effective treatment for metastatic cancer has remained elusive due to the persistence of drug-resistant metastasis stem cells (MetSCs) that drive relapse. MetSCs are tumor cell subpopulations enriched for their ability to reinitiate and sustain metastatic growth, displaying phenotypic plasticity and resistance to chemotherapy. These cells express the L1 cell adhesion molecule (L1CAM), a transmembrane protein detected in numerous human solid tumor types and at multiple disseminated organ sites. As a selective surface marker of MetSCs, L1CAM is a promising candidate for molecularly targeted drugs aimed at eliminating metastases, yet strategies to date have not achieved clinical success. Here, we develop antibody-drug conjugates to deliver highly toxic PNU-159682 payloads to L1CAM-expressing cells. We report the generation of monoclonal antibodies (mAb) with high binding affinity, specificity and selectivity for the human L1CAM extracellular domain. Optimized L1CAM-targeting mAbs were conjugated to PNU-159682 to generate ADC variants with both cleavable and non-cleavable linkers, with an average drug-antibody-ratio (DAR) of four. ADCs derived from three antibodies targeting various epitopes of the L1CAM extracellular portion potently killed cells exhibiting varying levels of surface L1CAM expression. L1CAM ADCs given as monotherapy resulted in robust tumor control and extended survival in mice harboring subcutaneous L1CAM^+^ xenografts or L1CAM^+^ lung metastases from triple-negative basal breast cancer and lung adenocarcinoma. Safety analyses with mouse cross-reactive antibodies indicate a feasible therapeutic window. Our findings offer strong proof-of-concept to support the preclinical development of these novel L1CAM ADCs as therapeutic agents for advanced solid tumors.

## Introduction

Metastasis is the principal cause of cancer mortality and the second-leading cause of death in the United States. A key challenge is that after surgery or systemic therapy, residual metastatic disease can persist as slow-cycling single cancer cells or cell clusters that are inherently resistant to anti-mitotic chemotherapy or to drugs targeting oncogenic drivers. Such metastasis stem cells (MetSC) ultimately reinitiate tumor regrowth and drive lethal relapse, highlighting the unmet clinical need to develop novel therapies that target this unique population of relapse-associated cancer cells (1–3). Overall, there remains a major unmet need for effective drugs that can address interrelated challenges in advanced solid tumors, namely to target disseminated lesions, overcome prior treatment resistance, and eradicate MetSC subpopulations in the primary tumor and at distant sites.

Recent work has identified the L1 cell adhesion molecule (L1CAM) as a marker of MetSC, with functional importance for the regeneration of aggressive tumors (4). Originally defined as a neuronal adhesion molecule (5,6), L1CAM is associated with metastasis and poor prognosis in several cancers, including colorectal, lung and breast (7–12). In mechanistic studies we demonstrated that L1CAM expression mediates cell spreading over capillaries to initiate proliferation in MetSC, and is critical for metastatic outgrowth in multiple organ sites such as the bone, lung, liver and brain (13,14). L1CAM^+^ MetSC remain closely associated with the vasculature in disseminated sites. In lung adenocarcinoma (LUAD) and colorectal cancer patient samples (4), L1CAM^+^ cells were strongly enriched in the residual disease that survived chemotherapy in primary tumors, and in metastases.

Antibody-drug conjugates (ADC) are a rapidly developing drug modality for cancer (15). ADCs are composed of a monoclonal antibody with target specificity, typically IgG1, conjugated to a drug payload via a chemical linker. By design, ADCs enable selective delivery of potent toxic payloads to cells expressing the target antigen, in turn mitigating systemic toxicities associated with free drug administration. The effectiveness and therapeutic potential of ADCs can be influenced by factors including immunogenicity and hydrophobicity of the molecule, and the efficiency of target recognition and internalization. As the field evolves, the toolbox for design of each ADC component continues to expand, with opportunities to optimize based on the class of toxic payload, linker release mechanism, conjugation technology, and mAb structure (15).

L1CAM is a strong candidate to pursue ADC development for multiple reasons. Although L1CAM-targeted antibodies have been developed and have demonstrated efficacy in tumor xenograft models, none has advanced to a clinical trial (16,17). This may be due in part to the fact that L1CAM has a large, multi-domain extracellular region interacting with numerous other receptors and components of the extracellular matrix (18). Antibodies binding to a specific epitope in the extracellular domain may not necessarily disrupt the specific interactions that allow L1CAM to promote metastatic outgrowth. On the other hand, as a cell adhesion molecule, L1CAM has a high intrinsic turnover rate, with continuous internalization. Therefore, an ADC approach can potentially overcome the limitations of prior anti-L1CAM agents by delivering a cytotoxic drug payload to MetSCs that express and internalize L1CAM. Furthermore, its relative lack of expression on non-tumor tissues suggests the opportunity for a therapeutic window to minimize off-target toxicities. Although L1CAM is expressed in certain neuronal and kidney tubular subpopulations, mouse studies suggest that L1CAM is essential during development, but there is no evidence of dependency in the adult brain.

Here we develop ADCs directed at human L1CAM, which display robust binding, cytotoxicity, and favorable safety and efficacy against L1CAM-expressing tumors *in vivo*. These represent promising agents for further clinical translation to target L1CAM-expressing treatment refractory and metastatic solid tumors.

## Materials and Methods

### Mouse models

All animal experiments were performed in accordance with protocols approved by the Memorial Sloan Kettering Cancer Center Institutional Animal Care and Use Committee (IACUC). Balb/c and A/J mice were procured from The Jackson Laboratory. AlivaMab© mice were sourced from Ablexis, LLC. For *in vivo* ADC experiments, NOD scid gamma (NSG) mice (Cat# 005557, Jackson Laboratory) aged 4 to 8 weeks were used, monitored for up to 30 weeks, and euthanized in accordance with IACUC protocols. Animals were maintained at temperatures of 21.1–22.2 °C, 30–70% humidity, 10–15 fresh air exchanges hourly, and a 12:12 h light / dark cycle (lights were on from 06:00–18:00). When monitoring the growth of metastasis, mice are anesthetized with 2-3% isoflurane/O_2_ and subjected to bioluminescence imaging (BLI) after retro-orbital injection of D-luciferin (150 mg/kg). BLI was performed in an IVIS Spectrum Xenogen instrument (PerkinElmer) and data were analyzed using Living Image Software v.4.5 (PerkinElmer).

### Patient-derived xenografts (PDXs)

LUAD PDXs were generated from primary and metastatic tumor samples as described (19). Briefly, samples were subcutaneously injected in the flank of a 6–8-week-old female mouse (Cat# 005557, Jackson Laboratory). Tumors were harvested when the volume reached 1500 mm^3^. Maximal tumor size was not exceeded. Serial passage was done by disaggregating and finely mincing the tumor, vigorously triturating in cold PBS, and straining through a 60 μm filter. Aliquots were cryopreserved in RPMI medium containing 10% DMSO or proceeded to generate PDX tumoroids as described below.

### Cell culture

MCF-7, MDA-MB-231, HEK293T cells were maintained in DMEM with 10% FBS and 100 units/mL penicillin/streptomycin. H23 and H524 cells were purchased from American Type Culture Collection (ATCC). H23, H524, and H2030 were cultured in RPMI with 10% FBS and 100 units/mL of penicillin/streptomycin. All cell lines were cultured at 37°C in a humidified incubator with 5% CO_2_. Prior to use, all cells tested negative for mycoplasma using the MycoAlert-PlusTM kit (Cat# LT07-710, Lonza). MDA-MB-231 LM2 cells were grown in DMEM with 10% FBS and 100 units/mL penicillin/streptomycin and resuspended in PBS for *in vivo* injection. Expi293F cells for antibody production were sourced from Thermo Fisher Scientific (A14527).

PDX tumoroids were cultured as described (13). Briefly, single cells were resuspended in culture media (RPMI with 10% FBS and 100 units/mL of penicillin/streptomycin) and mixed with chilled Matrigel (Corning Cat# 356231) at 1:1 ratio before placement in a 35-mm glass-bottom dish (MatTek Cat# P35G-1.5-14-C). Cells were seeded on top of the Matrigel-coated glass-bottom area of dishes at 5 x 10^4^ cells in cold, diluted Matrigel solution. The mixture was incubated at 37°C for 30 min to promote gelation, and additional culture medium was added gently to the side of the cell-containing Matrigel domes. Tumoroids were harvested in the Cell Recovery Solution (Corning Cat# 354270) for at least 30 min at 4°C on a rotator to digest Matrigel, and dissociated into single cells using TrypLE for 15 min at 37°C. Samples were strained through a 35 μm filter to remove cell clusters and counted before additional experiments.

### Immunohistochemistry of patient tissue sections

Patient samples were selected based on the availability of a sufficient quantity of formalin-fixed paraffin-embedded tissue from patients who had given preoperative consent to tissue utilization for research purposes. All human tissues were obtained under Memorial Sloan Kettering Institutional Review Board biospecimen research protocol 15-101. All patients provided pre-procedure informed consent. Tissue was generally processed within 1 h of surgical resection. Archival formalin-fixed, paraffin-embedded (FFPE) clinical tissue blocks for immunostaining were identified by database search and chart review. For immunohistochemistry (IHC) staining, tissue sections were processed and stained with anti-L1CAM antibody (Cat# 826701, BioLegend) via standardized automated protocols using a Ventana Discovery ST instrument. ImmPRESS horseradish peroxidase (HRP) anti-mouse IgG and ImmPACT DAB Peroxidase (Vector Labs) were used for detection. The slides were counterstained with hematoxylin, dehydrated and capped with a coverslip.

The IHC stained sample sections were digitally scanned using the Aperio AT2 slide scanner (Leica Biosystems) under 20x objective magnification. Scanned images of tissue sections were annotated for tumor compartments by pathologists. The membraneous or nuclear algorithm of Aperio ImageScope v12.4.3.7005 (Leica biosystems) was used under supervision of a pathologist to quantify the staining intensity of L1CAM and Sox2: 0+ (no staining), 1+ (weak staining), 2+ (moderate staining) or 3+ (strong staining). An H-score was assigned based on the intensity score and the percentage of scored cells: H-score = 0 x (% of cells with 0+ intensity score) + 1 x (% of cells with 1+ intensity score) + 2 x (% of cells with 2+ intensity score) + 3 x (% of cells with 3+ intensity score).

### Generation of L1CAM mAbs

Recombinant L1CAM proteins for immunizations and hybridoma screening were sourced commercially. The extracellular domain of human L1CAM (isoform 1, residues 1-1120) fused to a poly-histidine tag via its C-terminus was obtained from Creative Biomart (L1CAM-His).The extracellular portions of human L1CAM (isoform 1, residues 20-1120 with a heterologous signal sequence at the amino terminus) or mouse L1CAM (residues 1-1123), each with a human Fc fused to the C-terminus (L1CAM-Fc), were procured from R&D Systems. Cynomolgus macaque L1CAM-His was produced in-house with N- and C-terminal boundaries of isoform 3 matched with human L1CAM. All recombinant proteins were produced in mammalian cells.

Two cohorts of 6 mice each (Balb/c and A/J strains) and 8 AlivaMab® mice with a human immunoglobulin repertoire were primed with a mixture of human and mouse L1CAM-Fc, followed by 3 booster injections with human L1CAM-His over the course of 8-10 weeks. To further stimulate antigen presentation, 3 mice in each group were given agonistic anti-mouse CD40 antibody (FGK5 clone, BioXcell) on the day of immunogen administration and 3 days after. TiterMax® Gold (Sigma-Aldrich) was used as adjuvant for all doses. Immunogens were administered subcutaneously. Serum was collected periodically to assess the immune response to L1CAM by enzyme-linked immunosorbent assay (ELISA) using recombinant human L1CAM-His. Mice with high serum mid-point titers (1:10,000 or greater) after the third dose (primer plus 2 boosts) from each strain were selected for an acute boost with human L1CAM-His. Three days later, splenocytes were collected and fused with mouse myeloma cells using polyethylene glycol to generate hybridoma.

### Hybridoma screening

Hybridoma supernatants were analyzed using a screening funnel shown in Figure 2. The primary screen was carried out by ELISA using soluble, His-tagged human L1CAM and by flow cytometry using L1CAM-HEK293T cells. About 12% of the hits were cross-reactive with mouse L1CAM expressed by transiently transfected HEK293FT cells. Hybridoma were subcloned by limiting dilution, and clonal supernatants were tested for binding to human L1CAM by ELISA and flow cytometry. A subset of antibodies with a strong signal over background in the binding assays (ELISA, flow cytometry) was purified from hybridoma supernatant to test for binding to MDA-MB-231 cells. The top binders were tested for internalization in MDA-MB-231 cells, leading to a final list of antibodies that fulfilled the criteria defined by the screening funnel.

### Cross-reactivity, selectivity, and internalization of L1CAM mAbs

Candidate mAbs were purified from the supernatant of hybridoma clones for BLI analysis of binding to soluble human and cynomolgus macaque L1CAM-His. To narrow down the binding region of the development candidate mAbs, truncated versions of human L1CAM protein (Ig-like domains, residues 21-605; fibronectin-like type III (FNIII) repeats, residues 606-1116; each with a C-terminal 6x His tag) were recombinantly produced, followed by a BLI experiment to test binding at two concentrations (50 nM and 100 nM). A competition experiment was carried out within sub-domain proteins to identify epitope bins (Supplementary Table 1).

Binding kinetics with human L1CAM and ortholog species was measured by BLI using an Octet® Red96e instrument (ForteBio). His-tagged, monomeric human, mouse, or cynomolgus L1CAM protein was captured on Ni-NTA biosensors, and an 8-point, 2-fold dilution series of 143G03, 14A10 or 13F04 mAbs starting at 50 nM was used as analyte. The data was prepared by reference well subtraction and analyzed using ForteBio Data Analysis Package. Response curves were globally fit to a 1:1 Langmuir binding model.

To assess selectivity of the mAbs, ELISA plates were coated with various L1 family of cell adhesion molecules: human CHL-1 (residues 25-1096 with a C-terminal 10xHis tag, R&D Systems), human neurofascin (residues 25-1039 with a C-terminal 10xHis tag, R&D Systems) or human neuronal cell adhesion molecule (residues 30-600 with a C-terminal human IgG1 Fc tag, R&D Systems). Antibodies were added at various concentrations and positive control antibodies were included for each homolog. To analyze the internalization potency of antibodies, a Fab-ZAP assay was used. Briefly, saporin-conjugated Fab fragments binding to the Fc portion of human IgG1 were pre-incubated with L1CAM candidate mAbs or mouse IgG1 isotype control. The complex was added to cultures of L1CAM-HEK293T cells. Five days later, cell viability was analyzed using a colorimetric assay.

### Humanization of lead candidate mAbs

To humanize the parental murine 143G03 mAb, an approach utilizing a 3D structure generated by homology modeling and complementarity-determining regions (CDR) grafting was employed. A panel of 30 humanized antibody variants were generated from candidate sequences designed to maximize the amount of human sequence while retaining the parental mAb specificity and affinity. Selection of the candidate humanized mAb from the panel was determined based on evaluation of productivity in a mammalian expression system and binding to the target antigen by BLI and flow cytometry. The resultant candidate antibodies (the humanized 143G03, the AlivaMab® mouse-derived fully human 14A10, and the murine 13F04) were produced in stable CHO pools.

### Generation of L1CAM ADCs

The lead candidate mAbs 143G03, 14A10, and 13F04 were produced recombinantly in CHO cells. ADCs were produced in partnership with Abzena (Cambridge, UK) using the ThioBridge® platform. For ADC scouting work, small amounts of 143G03 and 14A10 mAbs were treated with tris (2-carboxyethyl)-phosphine (TCEP) for reduction of the interchain disulfides, followed by an addition of ThioBridge® linker-payload reagent. The mixture was incubated for varying timepoints and reaction was monitored by analytical SEC, HIC and LCMS for the optimal conjugation conditions. In initial screening experiments, six payloads with various mechanisms of action were selected for conjugation to 143G03 and 14A10, using a linker containing a valine-citrulline (Val-Cit) cathepsin B cleavage site, and a cyclic or linear PEG spacer unit. A matching isotype control IgG antibody was included in all reactions. ADCs were quantified by UV/VIS and characterized by various analytical methods to determine DAR rations (LC-MS), purity (SEC and SDS-PAGE) and endotoxin levels.

To generate the final 143G03 and 14A10 ADC molecules, cleavable and non-cleavable linkers were used. In parallel, 13F04 ADCwas generated using a non-cleavable linker. Analysis of the ADCs in *in vitro* mouse serum showed good linker stability across the different variants, with a reduction in average DAR of 3% or less.

### CRISPR-mediated knockout and shRNA knockdown of L1CAM

KN2.0 CRISPR, non-homology mediated, knockout (KO) kits were used to generate L1CAM KO cells (Origene Cat# KN411601) according to the manufacturer’s instruction. The constructs were transfected into cells using TurboFectin (Cat# TF81001, Origene), and cells were passaged for 2 weeks before isolating and expanding GFP^+^ single-cell knockout clones. Gene knockouts were validated by western immunoblotting of cell lysates. For knocking down L1CAM in PDXs, L1CAM human shRNA plasmid kit (Origene Cat# TL303597) was used. ShControl cells were generated in both cases by transducing the cells with scrambled shRNA control (Origene Cat# TR30021). Viral transductions were performed as stated below, and stable cell lines were generated by puromycin selection for 2 weeks.

### Viral transductions

Lentivirus suspensions were generated by transfecting 70-80% confluent HEK293T cells with lentiviral vectors using the lentiviral packaging kit (Cat# TR30037, Origene) or the Trans-Lentiviral shRNA packaging kit (Cat# TLP5912, Horizon Discovery) according to the manufacturer’s instruction. Alternatively, lentivirus particles were produced from HEK293T cells with psPAX2 and pMD2.G (Addgene plasmids 12260 &12259) using Lipofectamine 3000. Viral particles were collected, filtered through 0.45 μm sterile filters, and incubated with cells of interest for 24 h with 8 μg/mL of polybrene before recovering cells in complete media. Lentivirus particles were concentrated using Amicon Ultra-15 centrifugal filter unit according to the manufacturer’s instructions.

PDX-derived tumoroids were transduced using TransDux MAX (System Biosciences Cat# LV860A-1) based on the manufacturer’s instruction. Briefly, tumoroids were isolated from Matrigel and dissociated into single cells as described above. Single cells were resuspended in RPMI media mixed with MAX Enhancer and TransDux solutions with 8 mM HEPES. After adding lentivirus media, cells were transferred into a 24-well plate and spin-infected at 1500 g for 2 h at 32°C. After spinoculation, cells were resuspended in RPMI and Matrigel 1:1 solution and seeded back into a Matrigel-coated 35-mm glass-bottomed dish as tumoroid culture.

### Single-cell RNA sequencing

LUAD PDX tumoroids were cultured in Matrigel as described above, collected and dissociated into single cells prior to sequencing. Cells were sorted in ice-cold PBS with 0.04% BSA by flow cytometry using DAPI as a negative viability marker. Single-cell RNA-seq was performed on 10x Genomics Chromium platform. The scRNA-seq libraries were prepared based on the manufacturer’s protocol for 3’ end reading. The individual transcriptomes of encapsulated cells were barcoded during RT step and resulting cDNA purified with DynaBeads followed by amplification per manual guidelines. PCR-amplified product was fragmented, A-tailed, purified with 1.2X SPRI beads, ligated to the sequencing adapters and indexed by PCR. The indexed DNA libraries were double-size purified (0.6–0.8X) with SPRI beads and sequenced on Illumina NovaSeq S4 platform (R1 – 26 cycles, i7 – 8 cycles, R2 – 70 cycles or higher). The raw sequencing data were demultiplexed and aligned to the mouse genome using 10X CellRanger pipeline. This generated an output matrix of mRNA counts per cell for every gene. We used the filtered matrix for all downstream analysis.

All processing of the resulting data was performed using the Scanpy package (20). For quality control, cells with less than 200 genes detected and genes expressed by less than 3 cells were filtered out. Based on the library distribution, cells with less than 1,500 genes by counts were excluded from analysis. The total of 4,017 cells passed the filters.

The data were normalized by library size followed by a log transformation (base = 2, pseudo count = 1). From the normalized count matrices, principal component analysis (PCA) was performed using the top 4,000 highly variable genes (HVGs). The selected principal components (PCs) were used to construct k-nearest neighbor graph (n_neignbors = 30) to generate uniform manifold approximation and projection (UMAP) layouts, from which L1CAM expression was displayed.

### Flow Cytometry

HEK293T cells overexpressing human L1CAM as well as the MCF-7 and MDA-MB-231 human breast cancer cell lines were used. To assess the binding of L1CAM antibodies to cell surfaceexpressed L1CAM, antibodies were added to cells starting at 100 μg/mL, with 10 serial 3-fold dilutions, followed by washing in PBS. Cells were then incubated with an AlexaFluor 647 goat anti-human Fc secondary antibody. After washing, the stained cells were analyzed using a FACS Aria II SORP flow cytometer (Beckton Dickinson) or FACSymphony S6 high-speed cell sorter (Beckton Dickinson). The binding curves were calculated for each antibody and cell line normalized to secondary antibody only staining (MFI ratio).

### *In vitro* cytotoxicity assay

In pilot experiments, ADCs generated using 143G03 and 14A10 and one of six payloads were added to L1CAM-HEK293T, MCF-7, or MDA-MB-231 cells at various concentrations ranging from 0.1 pM to 100 nM, and cell viability was assessed 96 h later. ADCs generated using 143G03 and 14A10 plus a human IgG1 isotype control, conjugated to PNU-159682, with either a cleavable or a non-cleavable linker, were tested for cytotoxicity using L1CAM-HEK293T, MCF-7 and MDA-MB-231 cell cultures. The ADCs were added to the cells in 10 serial 5-fold dilutions starting at 10 μg/mL. Cell viability was assessed 96 h later using Cell-Titer-Glo (Cat# G9242, Promega), and cell viability was plotted against antibody concentration.

For testing across various cancer types, L1CAM ADC was generated using 13F04 conjugated to PNU-159682 with the non-cleavable linker. For treatment of cell cultures with ADCs, 2000 cells were seeded in 50 μL of culture media per well in the inner 60 wells of a white-bottom 96-well plate. PBS was added to the outer 36 wells to prevent evaporation in the test wells. After 24 h, ADC (13F04 or IgG1-ADC) in 50 μL of media were added to reach the indicated final concentrations. Plates were incubated for 3 days. 100 μL of Cell-Titer-Glo (Cat# G9242, Promega) was then added to each well and incubated at room temperature on a shaker for 10 min and plate was read in a microplate plate reader (BioTek, Synergy H1) to assess the cell viability. A microplate plate reader (BioTek, Synergy H1) was used to analyze Cell Titer Glo expression. Relative viability was quantified by normalization to controls that did not receive ADC.

For treatment of LUAD PDX tumoroids with ADCs, 2000 cells were pre-mixed with cold 12% Matrigel and ADC (13F04 or IgG1-ADC) and placed in wells of a 96-well plate, each well pre-coated with 25 μL of Matrigel. After gelation at 37°C, 100 μL of media was added on top of each sample. Plates were incubated for 5 days with ADCs or control. 100 μL of Cell-Titer-Glo (Cat# G9242, Promega) was then added to each well and incubated at room temperature on a shaker for 30 min. Cell viability was determined and quantified as described above.

### *In vivo* tumor models

For *in vivo* safety studies, ADCs or IgG1 control with cleavable and non-cleavable linkers were administered to female non-tumor bearing treatment-naïve athymic mice by intraperitoneal injection once a week, with four doses total. Mice were monitored weekly for weight and other clinical signs. Mice were sacrificed at the experimental endpoint and blood was collected for clinical chemistry analysis if needed.

For subcutaneous tumors, 5 x 10^6^ MDA-MB-231 cells were injected into the flank of athymic mice, and tumor growth was monitored by a digital caliper until they reached 100 mm^3^ in size. Four weekly doses of PNU-159682-conjugated 143G03 or 14A10 ADC with both cleavable (0.3 mg/kg) and non-cleavable linkers (1 mg/kg), as well as the equal amounts of IgG1 control ADC and vehicle control were administered intraperitoneally. Tumor volume was monitored weekly by using a digital caliper.

For lung colonization assays, 5 x 10^4^ MDA-MB-231 LM2 cells were introduced via tail vein injection into athymic mice to generate orthotopic lung metastases. Four weekly doses of PNU-159682-conjugated 143G03 or 14A10 ADC with both cleavable (0.3 mg/kg) and non-cleavable linkers (1 mg/kg), as well as IgG1 controls ADC and vehicle control were administered intraperitoneally. Tumor volume was monitored weekly by measurement of BLI as described above.

For treatment of LUAD metastasis-bearing mice with anti-L1CAM ADCs, NSG mice were inoculated with Ru323 or Ru631 LUAD PDX tumoroid cells (10^5^ cells per mouse) via the lateral tail vein. After metastatic engraftment (after 4 to 5 weeks), mice were separated into two groups with similar BLI intensities and treated with either IgG control ADC or anti-L1CAM ADC (13F04) in PBS, at 1 mg/kg, via intraperitoneal injection. For accessing the acute effect of ADC treatment, lung tissues were collected one week after a single dose of ADC. For testing the treatment efficacy and survival benefit, four doses of ADCs were administrated at weekly intervals. Tumor burden was tracked by BLI once or twice weekly throughout the course of experiment. Mice were euthanized once terminally ill, as mandated by IACUC.

### Western blotting

Protein lysates were prepared in ice-cold RIPA buffer (Sigma-Aldrich Cat# R0278) supplemented with Halt protease and phosphatase inhibitor cocktail (Thermo Fisher Scientific Cat# 78442). Cells were ruptured by sonication, and the lysates were centrifuged and pelleted at 4°C. The supernatant was collected and quantified using the BCA protein assay (Thermo Fisher Scientific Cat# 23228) analyzed on a microplate reader (BioTek Synergy-H1) at 562-nm absorbance with Gen5 software (BioTek). Samples were diluted in reducing Laemmli SDS sample buffer (Thermo Fisher Scientific, Cat# J60015.AC) and boiled for 5 min. Proteins were separated on NuPAGE Novex 4-12% Bis-Tris gels (Thermo Fisher Scientific Cat# NP0322BOX) using 1x MOPS SDS running buffer (Thermo Fisher Scientific Cat# NP0001) and transferred to nitrocellulose membranes. Once membranes were blocked with Intercept Tris-buffered saline (TBS) blocking buffer (LICOR Biosciences Cat# 927-60001) for 1 h at room temperature, they were probed with antibodies against β-actin (Sigma-Aldrich Cat# A1978, RRID: AB_476692) and L1CAM (BioLegend Cat# 826701, RRID: AB_2564904) in blocking buffer for overnight at 4°C. After washing three times with TBS with 0.1% Tween (TBST), the membranes were probed with secondary antibodies (LICOR Biosciences) in Intercept TBS blocking buffer for 1 h at room temperature. After another wash, bands were detected with the 680 and 800 channels of an Odyssey CLx imager (LICOR Biosciences).

### Immunofluorescence and fluorescence microscopy

Unstained slide sections were deparaffinized using Histo-Clear (National Diagnostics Cat# HS-200) and rehydrated in a series of ethanol washes. Antigen retrieval was performed in the Tris-based antigen unmasking solution (Vector Laboratories Cat# H-3300) in a steamer for 30 min. Sections were blocked in 10% normal goat serum (Life Technologies Cat# 50062Z) for at least 1 h at room temperature, and incubated with primary antibodies diluted in blocking solution overnight at 4°C. Antibodies used include mouse L1CAM (Miltenyi Biotec Cat# 130-115-812, AB_2727206), human L1CAM (Santa Cruz Biotechnology Cat# sc-53386, RRID: AB_628937), Sox2 (Invitrogen Cat# 14-9811-82, RRID: AB_891383), GFP (Aves Labs Cat# GFP-1010, RRID: AB_2307313), Cleaved Caspase-3 (Cell Signaling Technology Cat# 9661, RRID: AB_2341188), and human cytokeratin (Dako Cat# M3515, RRID: AB_2132885). After washing in PBS three times, samples were incubated with fluorophore-conjugated secondary antibodies for 1 h at room temperature in the dark, followed by another round of PBS washes and staining of nuclei with Hoechst 33342 (Thermo Fisher Scientific Cat# H3570). Sections were washed with PBS three times and mounted using ProLong diamond antifade mountant (Life Technologies Cat# P36970) before performing microscopy.

For fluorescence microscopy, slides were analyzed on a STELLARIS 8 confocal microscope (Leica), equipped with an Apo 63x 1.4 NA objective. Images were recorded with a Leica HyD (Leica Hybrid Detectors) controlled by Leica Application Suite X 4.5.0.25531 software. For scanning whole stained tissue sections, slides were scanned on a Pannoramic Scanner (3DHistech, Budapest, Hungary) using a 20x/0.8NA objective. Scans were visualized using Slideviewer (Version 2.7, 3DHistech) or ImageJ 1.53n.

### Bioinformatics analysis

Differential L1CAM expression across various tissue types was analyzed using gene chip– based data from the tumor and metastasis analysis pipeline of TNMplot (21). The source data include manually curated gene chip datasets of NCBI Gene Expression Omnibus (GEO). Patient survival data were curated from KM-plotter which integrates datasets from GEO, EGA, and TCGA (22). Overall survival was compared based on L1CAM mRNA expression levels, derived from either gene chip or RNA-seq data, with expression stratified into high and low groups using an automated best threshold cutoff. When profiling the LUAD PDX or cell line cohorts, samples were ranked according to their relative L1CAM expression levels or H-scores.

### Statistical analysis

An unpaired two-tailed Mann–Whitney test was used for two-group comparisons. All statistical analyses were performed using GraphPad Prism v.10.2.3 for Mac (GraphPad Software, www.graphpad.com). All bar graphs show mean values with error bars representing either S.E.M. or S.D., as defined in the figure legend. P values are indicated in the figure legends and, when appropriate, were rounded to the nearest single digit. P values of less than 0.05 were considered significant.

### Data Availability

The data generated in this study are available within the article and its supplementary files. The sequencing data have been submitted to the Gene Expression Omnibus under accession number GSE307811. Any additional information and related data are available from the corresponding authors upon reasonable request.

## Results

### Therapeutic potential of L1CAM across cancer types

To define the suitability of L1CAM as a therapeutic target for advanced solid tumors, we evaluated its expression in patient samples using integrated datasets from TCGA, EGA, and GEO (21,22). A pan-cancer survey revealed that L1CAM levels were elevated in many metastatic tumors compared to primary tumors, and that high L1CAM expression correlated with decreased overall survival across multiple carcinoma types, including in lung adenocarcinoma (LUAD) and basal (associated with triple negative) breast cancer (**Fig. 1A** and **B**). Interestingly, the combined analysis of expression differentiation and survival association by L1CAM was most pronounced in LUAD (**Fig. 1A** and **B**), which we subsequently selected as a focus for evaluating ADC efficacy. Consistent with this analysis, immunohistochemistry staining revealed robust L1CAM expression in LUAD metastases compared to primary tumor (**Fig. 1C**). We analyzed a cohort of LUAD patients from whom patient-derived xenografts (PDXs) were generated, revealing a range of L1CAM expression spanning clinicogenomic profiles (**Fig. 1D**). Notably, L1CAM expression was maintained in derived tumoroids and LUAD cell lines independent of KRAS mutation status (**Fig. 1E** and **F**). These data are consistent with published literature linking L1CAM expression to advanced disease across multiple tumor types (4,7–12) and support its relevance as a therapeutic target for delivering molecularly targeted therapy to metastatic stem cells within these lesions.

**Figure 1.**
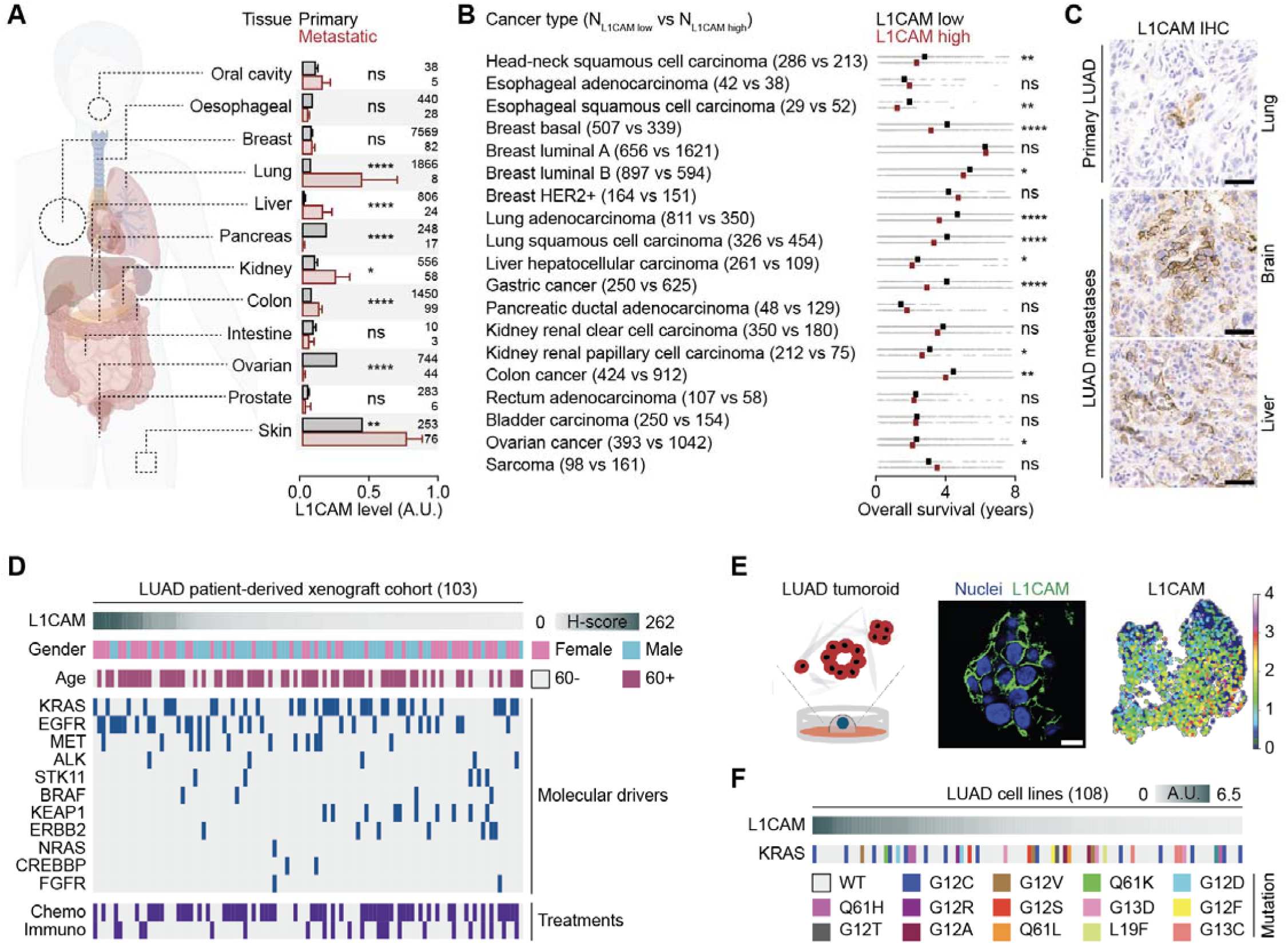
L1CAM as a prognostic marker and a therapeutic target in various cancer types. **A**, L1CAM expression in primary and metastatic tumors from different tissues. Bar graph, mean ± S.E.M. Sample sizes are indicated in the figure. **B**, the overall survival rate for various cancer types with respect to L1CAM expression. Overall survival, individual data points in grey; median survival in black and red. Sample sizes are indicated in the figure. **C**, L1CAM IHC staining of LUAD primary tumor and metastases from patient tissue sections. Scale bar, 50 µm. **D**, Clinical profiles of LUAD PDX cohort ranked by L1CAM H-score. Each column represents a single patient from whom the xenograft was derived. Sample sizes are indicated in the figure. **E**, Schematic representation of LUAD PDX-derived tumoroid culture, followed by immunofluorescence imaging of L1CAM staining and UMAP analysis of L1CAM transcript levels from scRNA-seq data (4,017 cells). Scale bar, 10 µm. **F**, Expression of L1CAM in various LUAD cell lines with respect to KRAS mutation status. Each column represents a single cell line. Sample sizes are indicated in the figure. A.U., arbitrary unit. Statistical significance was assessed using a two-tailed Mann-Whitney test (**A**,**B**). ns, not significant; \**P* < 0.05; \*\**P* < 0.01; \*\*\*\**P* < 0.0001.

### Generation of antibodies to the L1CAM extracellular domain

As the first step in our ADC design, we sought to generate antibodies with high-affinity binding to the extracellular region of human L1CAM (**Fig. 2A**). We began by immunizing mice with recombinant L1CAM proteins and screened hybridoma supernatants for antibodies that demonstrated *in vitro* binding to recombinant L1CAM protein and to L1CAM expressed by cells (**Fig. 2B**). Binding to soluble human L1CAM was determined by ELISA (**Fig. 2B**). Next, to determine binding to cell surface L1CAM, we performed flow cytometry using human embryonic kidney cells (HEK293T/FT cells) overexpressing human or mouse L1CAM (hereon, 293T-L1CAM) and two human breast cancer cell lines that endogenously express L1CAM: MCF-7 (ER/PR+, luminal A subtype) and MDA-MB-231 (ER/PR/HER2-, basal subtype).

**Figure 2.**
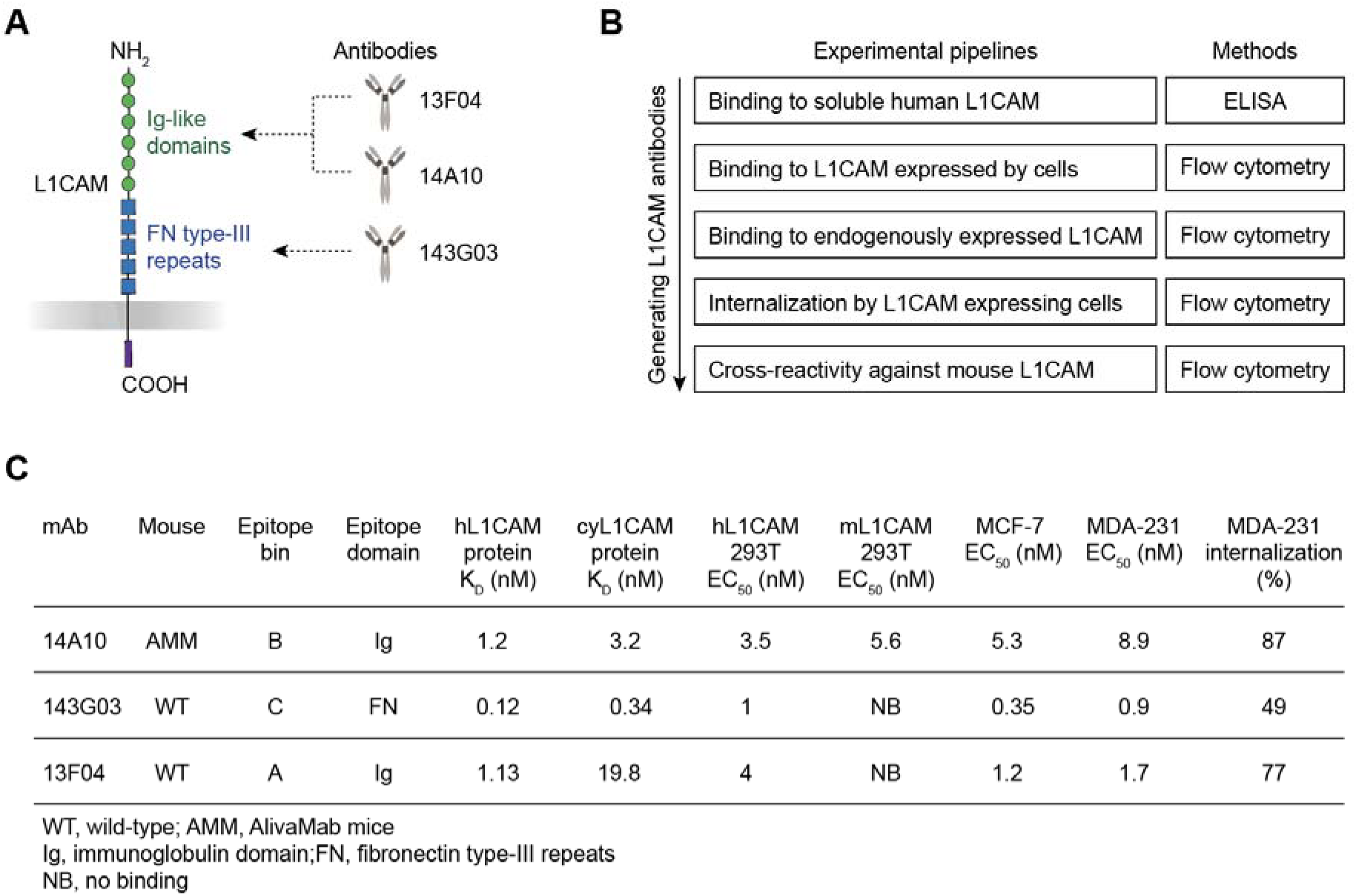
Generation of L1CAM antibodies binding to the extracellular portion. **A**, Schematic representation of the L1CAM molecule and its extracellular domains as binding sites for the generated antibodies. **B**, Workflow for hit selection of hybridoma supernatants from mice immunized with L1CAM immunogens. **C**, Molecular characterization of the L1CAM antibodies derived from the experimental steps outlined in panel **B**.

After hybridoma subcloning and sequencing, lead candidate mAbs were produced recombinantly. The EC_50_ values for binding to various cell types expressing L1CAM were in the single-digit nM or sub-nM range (**Fig. 2C**; Supplementary Table S1). To determine the potential for internalization, we performed a Fab-ZAP assay in 293T-L1CAM cell cultures. We found that all L1CAM lead candidate mAbs were extensively internalized after 5 days of incubation, after which their cross-reactivity to cynomolgus macaque and mouse His-tagged L1CAM was evaluated (**Fig. 2C**).

Based on the screening results, three candidate 143G03, 14A10, and 13F04 mAbs were prioritized for ADC formatting and testing. Using a BLI competition assay, we determined that 13F04 and 14A10 bind to the Ig-like domains, whereas 143G03 binds to the fibronectin-like type III (FNIII) repeats of the L1CAM extracellular domain (**Fig. 2A** and **C**). In addition, 14A10 is cross-reactive with mouse L1CAM, allowing for preclinical assessment of on-target toxicity in mice.

### Optimization of payloads for L1CAM ADCs

We utilized a disulfide re-bridging platform (ThioBridge™, Abzena) to conjugate cytotoxic payloads to the 143G03 and 14A10 L1CAM mAbs. This technology employs the selective reduction of disulfide bonds between the heavy and light chains as well as the hinge region at the heavy-heavy chain interface of the antibodies. Conjugation of a bis-sulfone cysteine-bridging linker under mild aqueous conditions results in the production of stable ADCs with highly homogeneous drug-to-antibody ratios (DAR) (23–25). We conducted initial experiments evaluating five payloads with various mechanisms of toxicity: microtubule polymerase inhibitors (monomethyl auristatin E [MMAE] and ABZ038), DNA alkylator and/or intercalators (Duocarmycin TM and PNU-159682), and RNA Polymerase II inhibitor (α-Amanitin). Each payload was linked with a valine-citrulline (Val-Cit) cathepsin B cleavage site for release of the payload in the lysosome, and a PEG spacer to reduce hydrophobicity of the ADCs. An isotype control IgG1 antibody was included in all reactions. All ADC variants had an average drug-to-antibody ratio (DAR) of 4.

We determined the potency of the ADCs in cytotoxicity assays using cell lines that display membrane-bound human L1CAM at different levels: stably transfected 293T-L1CAM cells (overexpression), MCF-7 (medium expression) and MDA-MB-231 (low-medium expression) human breast cancer cell lines. ADCs were added to cell cultures at concentrations ranging from 0.1 pM - 100 nM and cell viability was assessed after 96 h (**Fig. 3A**-**E**). Of the 6 payloads used, PNU-159682 exhibited robust killing of all three cell lines including MDA-MB-231 cells, which express lower endogenous levels of L1CAM (**Fig. 3A**). Therefore, PNU-159682, a metabolite of the anthracycline nemorubicin, was selected for further optimization and proof-of-concept evaluation in animal models.

**Figure 3.**
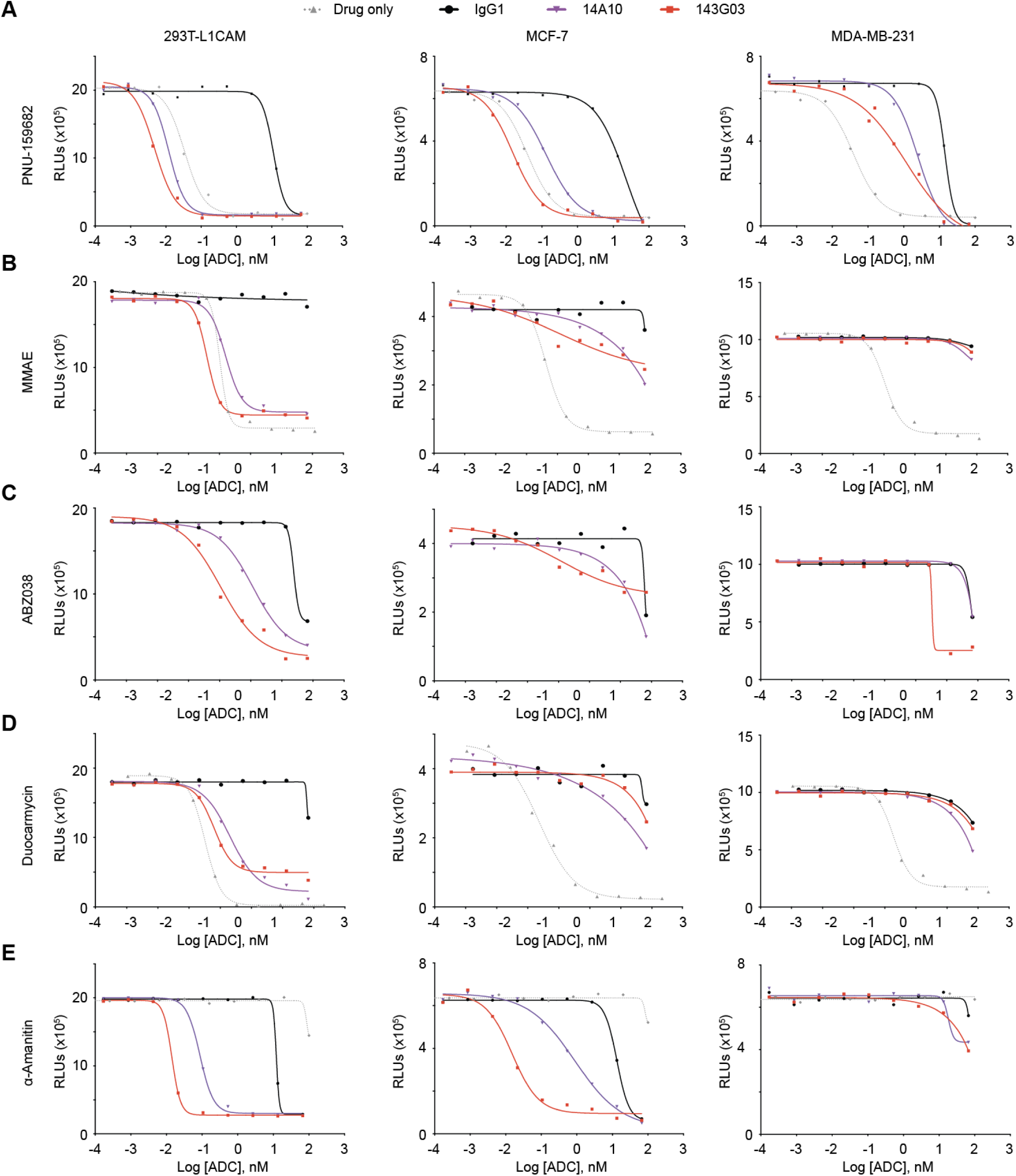
Payload optimization for L1CAM ADCs. Cytotoxicity curves for L1CAM ADCs generated from either the 14A10 (purple) or 143G03 (red) antibody conjugated to PNU-159682 (**A**), MMAE (**B**), ABZ038 (**C**), Duocarmycin (**D**), or α-Amanitin (**E**) using a Val-Cit linker. Cell viability was measured as a function of ADC concentration (0.1 pM to 100 nM) after 96 h incubation with HEK293T cells overexpressing L1CAM (293T-L1CAM) or MCF-7 and MDA-MB-231 human breast cancer cell lines. A mouse IgG1 isotype control antibody conjugated to the corresponding payload (black) and the corresponding free drug (dotted grey) were included as controls.

Of note, we observed killing potencies of 3- to 10-fold higher for ADCs derived from 143G03 compared to 14A10, which correlates with the respective binding affinities of the two antibodies for L1CAM (**Fig. 2C**). Non-specific killing was observed for the IgG1 isotype control at higher concentrations for several payloads (**Fig. 3A**-**E**), suggesting additional uptake mechanisms known to be associated with ADCs, including Fc receptor-mediated endocytosis or pinocytosis (26). The final lead candidate ADCs were further evaluated for affinity, solubility, stability and immunogenicity.

### Optimization of linkers for L1CAM ADCs

To optimize an ADC linker, humanized 143G03 and fully human 14A10 mAbs were conjugated to PNU-159682 using a cleavable linker (**Fig. 4A**) or using a non-cleavable tri-glycine linker (**Fig. 4B**). ADCs with cleavable linker and non-cleavable linkers were stable in mouse and human sera for 96 h, with less than 3% and no payload release, respectively, as detected by LC-MS.

**Figure 4.**
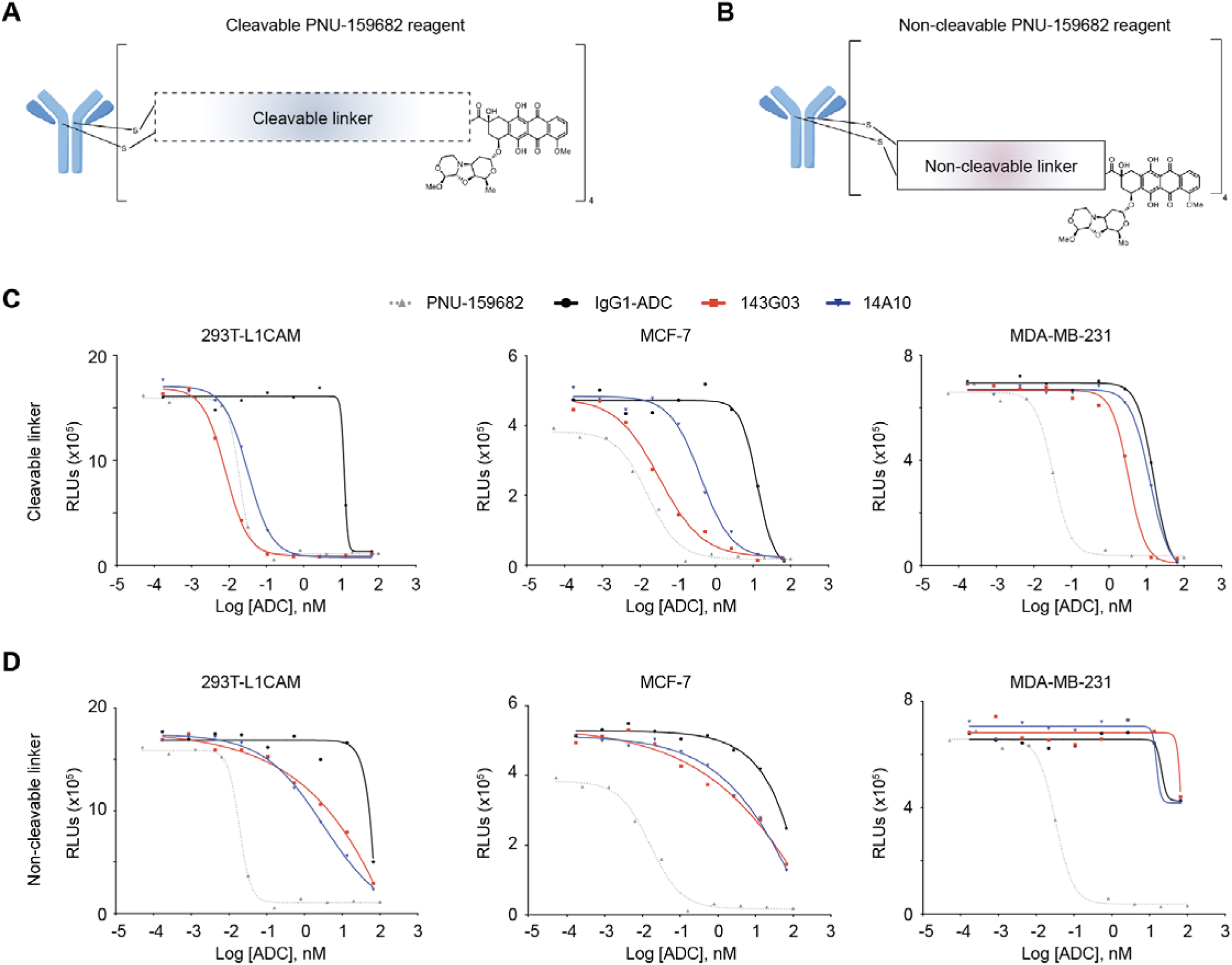
Linker optimization for L1CAM ADCs. Schematic representations of the ADC with cleavable (**A**) or non-cleavable (**B**) linkers and the PNU-159682 payload. Cytotoxicity curves for 143G03 (red) and 14A10 (blue) ADCs conjugated to PNU-159682 with either a cleavable (**C**) or non-cleavable (**D**) linker in 293T-L1CAM, MCF-7, and MDA-MB-231 cell lines. An ADC generated from a mouse IgG1 isotype control antibody (IgG1-ADC; black) and free PNU-159682 (dotted grey) were included as controls.

143G03 and 14A10 ADCs conjugated to PNU-159682 with cleavable linkers were potently cytotoxic when administered to 293T-L1CAM and MCF-7 cells, while the effects were more moderate in MDA-MB-231 cultures (**Fig. 4C**). ADCs with the non-cleavable linkers showed effective but limited potency *in vitro* (**Fig. 4D**), which is likely attributed to the requirement for degradation in the lysosome for payload release. Although the ADCs with cleavable linkers exhibited greater potency *in vitro*, both linker types demonstrated selective cytotoxicity in the sub-nanomolar range. Given that L1CAM expression and turnover may be different in an *in vivo* tumor setting, we opted to test the ADCs with both cleavable and non-cleavable linkers in subsequent *in vivo* experiments.

### *In vivo* safety and optimization of L1CAM ADC treatments

We initially assessed the safety of ADC treatments in mice. 143G03 and 14A10 ADCs were conjugated with either cleavable or non-cleavable linkers to the PNU-159682 payload and administered once weekly via intraperitoneal injection to treatment-naïve female NSG mice for a total of four weeks. Throughout the study, we monitored the mice for body weight and clinical signs (Supplementary Fig. S1 and S2). At the time of sacrifice, blood was collected for complete blood count (CBC) (Supplementary Fig. S1B-S1P) and clinical chemistry analysis (Supplementary Fig. S2A-S2Z). Mouse organs were collected, sectioned and stained for tissue-specific histological assessments of ADC treatments (Supplementary Table 2). Notably, histopathological evaluation of brain and peripheral tissues in animals treated with mouse cross-reactive ADC (14A10) by a trained veterinary pathologist revealed no evidence of perineural inflammation, apoptosis or other neuronal pathology.

Mice treated with 1 mg/kg of ADCs containing cleavable linkers experienced approximately 20% body weight loss within the first 10 days (Supplementary Fig. S1A). The dose was subsequently reduced to 0.5 mg/kg, at which point only minor weight loss and other clinical signs were observed after the third and fourth doses (Supplementary Fig. S1 and S2). For *in vivo* efficacy studies, the final dose of ADCs with cleavable linkers was set at 0.3 mg/kg. In contrast, ADCs with non-cleavable linkers were well tolerated at doses up to 1 mg/kg (Supplementary Fig. S1 and S2), which was selected as the final dose for efficacy testing.

To optimize *in vivo* anti-tumor activity of ADC treatments, we treated either subcutaneous MDA-MB-231 LM2 tumors from the flanks of athymic mice (**Fig. 5A** and **B**) or lung metastases of MDA-MB-231 LM2 from tail vein injection (**Fig. 5C** and **D**) with four weekly doses of 143G03 or 14A10 ADCs with cleavable or non-cleavable linkers. Subcutaneous tumors were allowed to grow to 100 mm^3^ in size, and lung metastasis burden was allowed to reach a bioluminescence (BLI) intensity of 10^4^ p/s/cm^2^/sr before the first ADC infusion. All L1CAM ADCs demonstrated sustained tumor control even after treatment withdrawal, resulting in improved overall survival (**Fig. 5A**-**D**). Strikingly, L1CAM ADCs with non-cleavable linkers induced complete tumor regression in both subcutaneous tumor and lung metastasis models, resulting in 100% long-term survival (**Fig. 5A**-**D**). Therefore, our findings indicate that ADCs with a non-cleavable linker to the PNU-159682 payload exhibit improved safety and superior *in vivo* anti-tumor efficacy.

**Figure 5.**
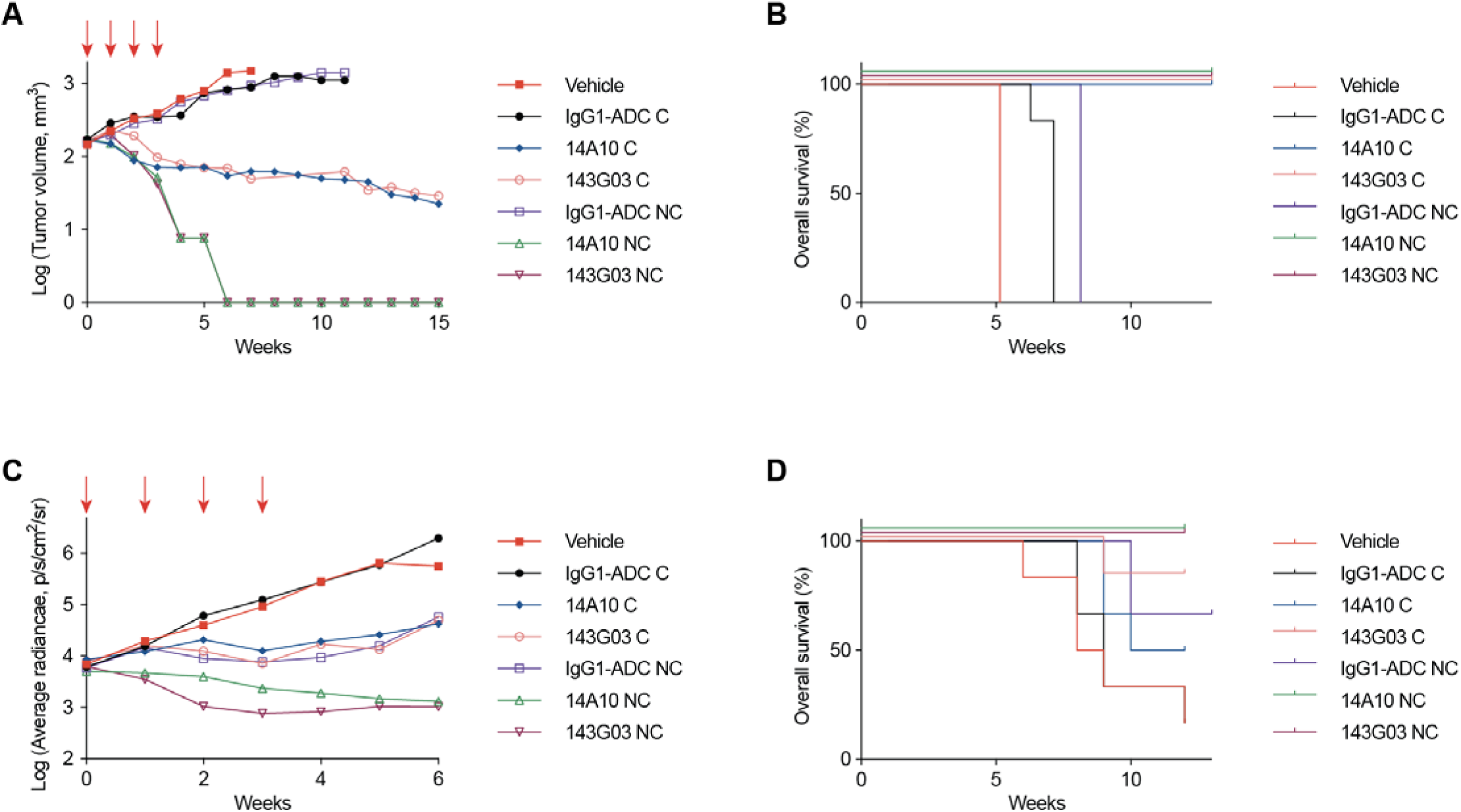
*In vivo* efficacy of L1CAM ADCs against Triple Negative Breast Cancer. Athymic mice were transplanted subcutaneously (**A**, **B**) or via tail vein injection (**C**, **D**) with MDA-MB-231LM2 breast cancer cells. Mice were administered 4 weekly doses (shown in red arrows) of ADCs generated with the 143G03 or 14A10 ADC conjugated to PNU-159682 with a cleavable (C) linker, (0.3 mg/kg) or non-cleavable (NC) linker (1 mg/kg), and IgG1-ADC or vehicle control. Tumor volume (**A**), tumor growth (**C**) and overall survival (**B**, **D**) were monitored on a weekly basis.

### Anti-tumor effect of L1CAM ADC across cancer types

Next, we generated the 13F04 ADC using the optimized combination of linker and payload described above. To assess its potency across various cancer types, cytotoxicity assays were performed on 293T-L1CAM cells, breast cancer cell lines (MCF-7 and MDA-MB-231), a small cell lung cancer line (H524), lung adenocarcinoma (LUAD) cell lines (H2030 and H23), and LUAD patient-derived xenograft (PDX) tumoroids (Ru631). Compared with an IgG1 control ADC, 13F04 ADC showed strong, selective cytotoxicity in the sub-nanomolar range against 293T-L1CAM, breast cancer and lung cancer cell lines (**Fig. 6A**-**F**). This cytotoxic effect was eliminated by knocking out L1CAM in H23 cells (**Fig. 6G**; Supplementary Fig. S3A). Additionally, 13F04 ADC effectively targeted L1CAM^+^ tumoroids generated from PDX Ru631 and this effect was abolished in L1CAM knockdown Ru631 tumoroids (**Fig. 6H** and **I**; Supplementary Fig. S3B).

**Figure 6.**
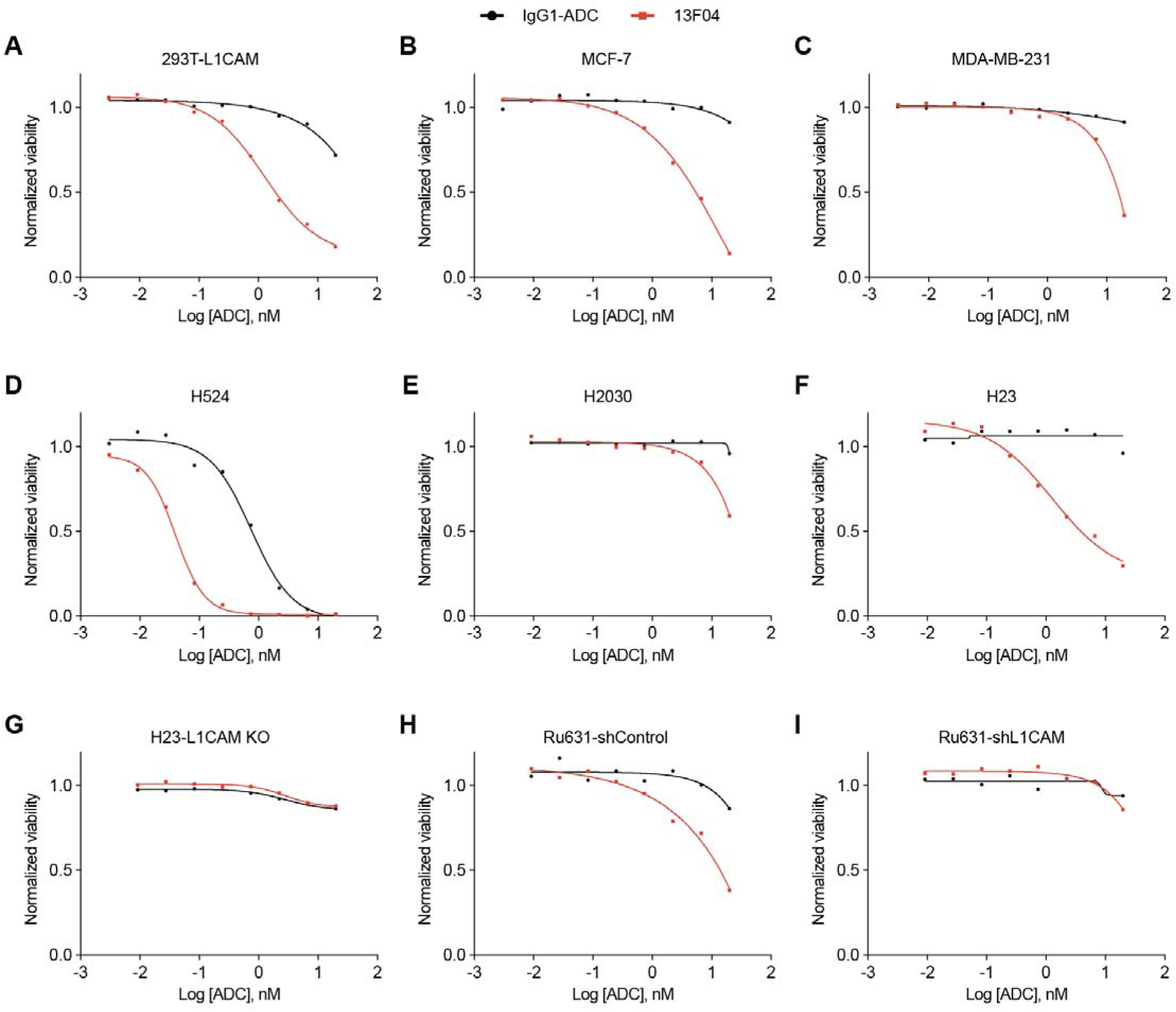
*In vitro* efficacy of L1CAM ADC across various cancer types. Cytotoxicity curves for 13F04 ADC (red) conjugated to PNU-159682 with a non-cleavable linker in the following cell lines: 293T-L1CAM (**A**), MCF-7 (**B**), MDA-MB-231 (**C**), H524 (**D**), H2030 (**E**), H23 parental (**F**), and H23 with L1CAM knockout (**G**). Efficacy was also evaluated in PDX-derived Ru631 tumoroids with either shControl (**H**) or shL1CAM knockdown (**I**). An ADC generated from a mouse IgG1 isotype control antibody (IgG1-ADC; black) was used as a negative control.

To assess the effectiveness of 13F04 ADC at targeting L1CAM^+^ metastatic cells, we used the Ru323 PDX model, which was derived from an untreated EGFR-mutant LUAD primary tumor, and the Ru631 PDX model, which was derived from an untreated KRAS-mutant LUAD spine metastasis (13). Approximately 60% of the cells in both PDX models are L1CAM^+^ (Supplementary Fig. S3C and S3D), making these PDXs suitable for testing the efficacy of 13F04 ADC against metastatic tumor growth. We first analyzed the acute response of Ru631 cells to treatment with 13F04 ADC. One week after intravenous inoculation of Ru631 cells into NSG mice, the mice were treated with a single dose (1 mg/kg) of 13F04 ADC or control ADC by intraperitoneal injection. One week later, the incipient Ru631 tumors from control ADC group showed a high proportion of L1CAM^+^ cancer cells and L1CAM staining at cell-cell interfaces, whereas Ru631 tumors from 13F04 ADC treated mice showed a markedly decreased proportion of L1CAM^+^ cells and the few that remained showed internalized L1CAM (**Fig. 7A** and **B**). Ru631 lung metastatic tumors that were allowed to grow for 4 weeks showed some enrichment of L1CAM^+^ cells during early metastatic outgrowth (Supplementary Fig. S3E and S3F). Treating with a single dose of 13F04 ADC showed a scattered histology (**Fig. 7C**), a reduction in tumor size (**Fig. 7D**), presence of apoptotic cells as determined by cleaved caspase 3 staining (**Fig. 7E** and **F**), and a marked decrease in the fraction of SOX2^+^ progenitor cells compared to counterparts treated with control ADC (**Fig. 7G** and **H**).

**Figure 7.**
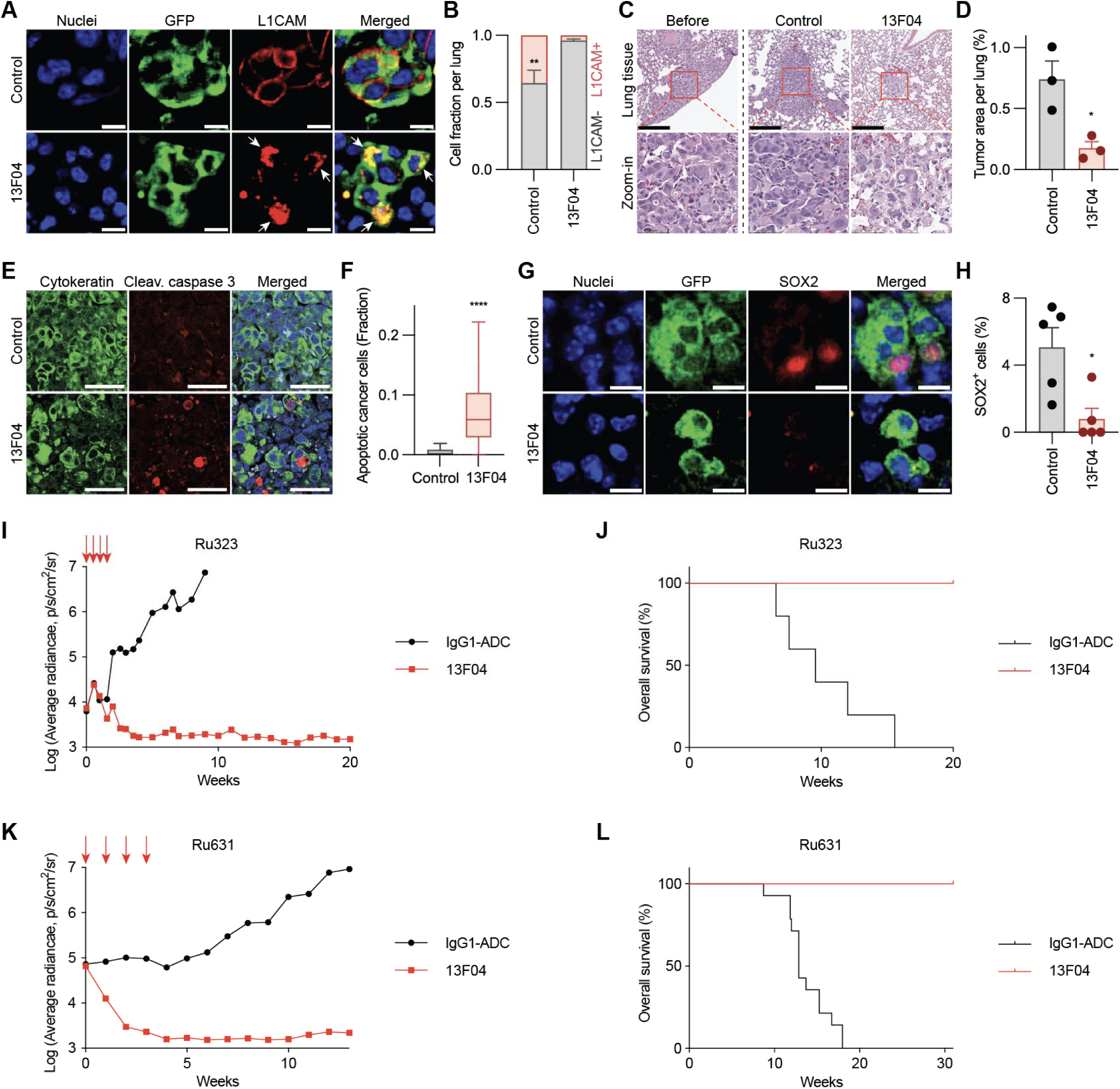
*In vivo* efficacy of L1CAM ADC against Lung adenocarcinoma (LUAD) metastasis. **A**, NSG mice were inoculated via the tail vein with GFP-expressing Ru631 LUAD PDX cells. One week later, mice were treated with a single dose of 13F04 ADC or IgG control ADC, and lungs were harvested and analyzed by IF staining for GFP (cancer cells) and L1CAM in one week after ADC treatment. White arrows, cytoplasmic L1CAM staining upon anti-L1CAM ADC treatment. Scale bar, 10 µm. **B**, Quantification of L1CAM^−^ or L1CAM^+^ fraction of Ru631 PDX treated with ADC in the experiment of panel (**A**). *n* = 3,195 cells (Control); 2,110 cells (13F04). \*\**P* = 0.0079. **C**, H&E staining of lung sections before or one week after a single dose of ADC treatment, administered four weeks after inoculation of Ru631 PDX cells into NSG mice via the tail vein. Magnified regions are shown (*red boxes*). Scale bar, 200 µm. **D**, Bar graph showing the percent area of metastatic lesions per lung in the experiments of panel (**C**). *n =* 3 per experimental condition. \**P* = 0.0239. **E**, IF staining for human pan-cytokeratin (cancer cells) and cleaved caspase 3 in lung metastatic colonies one week after a single dose of ADC treatment, administered four weeks post-inoculation of Ru631 PDX cancer cells into NSG mice via the tail vein. Scale bar, 50 µm. **F**, Fraction of cleaved caspase 3^+^ (apoptotic) cells detected by IF in the experiment of panel (**E**). *n* = 3,257 cells (Control); 788 cells (13F04). \*\*\*\**P* < 0.0001. **G**, IF staining for GFP (cancer cells) and SOX2 one week after a single dose of ADC treatment, administered one week after inoculation of Ru631 PDX cancer cells into NSG mice via the tail vein. Scale bar, 10 µm. **H**, Percentage of SOX2^+^ cancer cells detected by IF in the experiment of panel (**G**). *n* = 5 per experimental condition. \**P* = 0.0317. **I**-**L**, Therapeutic assessment of LUAD metastasis treated with L1CAM or IgG ADCs. Single-cell suspensions from LUAD PDX tumoroids transduced with luciferase were injected into NSG mice via the tail vein. After 4 to 5.5 weeks of metastatic engraftment and outgrowth in the lungs, the mice received control ADC (1 mg/kg) or 13F04 ADC (1 mg/kg) by intraperitoneal injection once weekly for four weeks at the indicated times (*red arrows*). Tumor burden was monitored by BLI weekly. **I**, Metastatic burden of mice inoculated with Ru323 PDX tumoroid cells and treated with control ADC or 13F04 anti-L1CAM ADC. *n =* 5 mice per group. \*\*\*\**P* < 0.0001. **J**, Overall survival plots of mice in the experiment of panel (**I**) \*\**P* = 0.0018. **K**, Metastatic burden of mice inoculated with Ru631 PDX tumoroid cells and treated with control ADC or 13F04 anti-L1CAM ADC. Control, *n =* 14 mice; 13F04, *n =* 15 mice. \*\*\*\**P* < 0.0001. **L**, Overall survival plots of mice in the experiment of panel (**K**). \*\*\*\**P* < 0.0001. The bar graph indicates mean ± S.E.M. (**B**,**D**,**H**). Data are shown as a box (median ± 25-75%) and whisker (maximum to minimum values) plot (**F**). Statistical significance was assessed using the two-tailed Mann-Whitney test (**B**,**F**,**H**), and two-tailed *t* test after passing the Shapiro-Wilk normality test (**D**).

To determine the efficacy of 13F04 ADC against established metastatic tumors, we performed intravenous inoculating of Ru323 or Ru631 cells in NSG mice and allowing metastatic growth in the lungs to become established over 4 to 5 weeks before treating the mice with four weekly doses (1 mg/kg) of control ADC or 13F04 ADC (**Fig. 7I**-**L**). Lung metastasis tumor burden was assessed by BLI of a luciferase vector expressed in the cancer cells. In the control ADC group, the metastatic burden grew exponentially (**Fig. 7I** and **K**) and the animals began to die within 11 weeks (**Fig. 7J** and **L**). In contrast, 13F04 ADC treatment resulted in a marked reduction of metastatic burden after two doses, with complete eradication of detectable tumors after four doses (**Fig. 7I** and **K**). The 13F04 ADC-treated mice survived for up to 9 months without signs of relapse, while all the mice in the control groups succumbed to the disease within 4 months (**Fig. 7J** and **L**). These results provided additional evidence that L1CAM^+^ cells are essential for initiating and sustaining metastatic growth in these cancer models, which can be effectively targeted using L1CAM ADCs.

## Discussion

We have shown a proof-of-concept study for the development of ADCs directed at L1CAM, as a strategy to target advanced and metastatic solid tumors. We generated monoclonal antibodies to human L1CAM, and demonstrated robust binding to endogenously expressed L1CAM and internalization by L1CAM-expressing cancer cells, with minimal cross-reactivity to related protein homologs. By conjugation to an anthracycline derivative payload, PNU-159682, the antibodies showed strong cytotoxic effects *in vitro*, and ability to control the growth of subcutaneous and metastatic lesions *in vivo*. Safety assessments indicate a suitable therapeutic window for L1CAM-directed therapy.

L1CAM ADC formatting was optimized using two lead antibody candidates, 14A10 and 143G03, which bind to the Ig-like domains and FNIII repeats of the L1CAM extracellular region, respectively. Regardless of the epitope, the affinity for L1CAM and the anti-tumor potency of ADCs using either antibody were encouraging. Among the five payloads tested, PNU-159682 exhibited the strongest *in vitro* cytotoxic effect particularly when conjugated via a cleavable linker. In contrast, ADCs with a non-cleavable linker demonstrated superior anti-tumor activity *in vivo*. This may be attributed to differences in safety and tolerability, which allowed for a higher administered dose of the non-cleavable format (1 mg/kg) compared to the cleavable format (0.3 mg/kg). Additionally, organ-specific factors may contribute to the relatively reduced efficacy of cleavable linker compounds within the lung tumor microenvironment.

Optimization of ADC design enabled the development of a third L1CAM ADC using the lead candidate antibody 13F04, which demonstrated the best combined binding and internalization efficiency in the MDA-MB-231 breast cancer cell line. The corresponding 13F04 ADC showed potent anti-tumor activity across multiple cancer types and effectively eliminated PDX-derived LUAD metastases *in vivo*. These findings support a strategy to eradicate metastatic progenitors via L1CAM-targeted intervention.

There is an unmet need for drugs capable of targeting and eliminating therapy-resistant disease in patients with advanced solid tumors. The elimination of quiescence-capable MetSC with an L1CAM ADC is expected to be complementary to chemotherapy and therapies that target rapidly proliferating cells. Furthermore, plasticity poses a great challenge for cancer treatment, since tumors may evade therapy by dynamically altering their phenotype. However, entry into a regenerative L1CAM^+^ MetSC state represents a conserved bottleneck of advanced cancers (4). This vulnerability represents a novel, promising concept in cancer therapeutics that addresses the plasticity of advanced cancer driving metastatic relapse after therapy. Initially, L1CAM ADCs may be used to treat residual disease in patients with treatment refractory advanced solid tumors. Eventually, L1CAM ADCs may also have utility in the adjuvant/neoadjuvant setting to treat latent micrometastatic disease and prevent lethal macrometastatic relapse.

## Acknowledgments

We thank the MSKCC Antitumor Assessment Core, Molecular Cytology Core Facility, Flow Cytometry Core Facility, Pathology Core Facility, and Animal Imaging Core for their technical assistance. We thank Y Kaygusuz for antibody internalization assays and J Simundza for assistance with the manuscript. We thank FE Kombak and M Jain for IHC analysis. This work was supported by National Institutes of Health grants R35-CA263816 (CMR), R35-CA252978 (JM), P01-CA129243 (JM), R37-CA266185 (KG), K08-CA230213 (KG), P30-CA008748 (MSKCC), grants from the Alan and Sandra Gerry Metastasis and Tumor Ecosystems Center at MSKCC (JM), Experimental Therapeutics Center (JM, KG), Therapeutics Discovery Fund (JM, KG), Burroughs Wellcome Career Awards for Medical Scientists (KG), and the Translational Research Oncology Training Program at MSKCC (JSP).

## Conflicts of Interest

JM, KG, AGK, SRK and ICL are inventors in U.S. Patent 11,464,874, and U.S. Provisional Patent Applications 63/478,809 and 63/478,829 on targeting L1CAM to treat cancer. JM has been a consultant and holds equity in Scholar Rock. KG has received honoraria from Genentech. CMR has consulted regarding oncology drug development with AbbVie, Amgen, AstraZeneca, Boehringer Ingelheim, Daiichi Sankyo, and Merck, and has received licensing and royalty fees for DLL3-directed therapies. CMR serves on the scientific advisory boards of Auron, DISCO, and Earli.

## Author Contributions

JSP, PJB, ICL, JM and KG conceived the study and designed the experiments; KG and LH performed cell culture and metastasis assays with 143G03 and 14A10 ADCs, and JSP and CK with 13F04 ADC; AGK, MAP, TEW, SRK, PJB, ICL, JM and KG developed and optimized L1CAM antibodies and ADCs; CMR generated and provided PDXs; JSP, JM, and KG wrote the manuscript. All authors edited and approved the manuscript.

## Tables of Supplemental Content

Supplementary Table 1: Binding profile of candidate L1CAM mAbs

Supplementary Table 2: Histological assessments of ADC-treated tissues

---

**Supplementary Figure S1.**
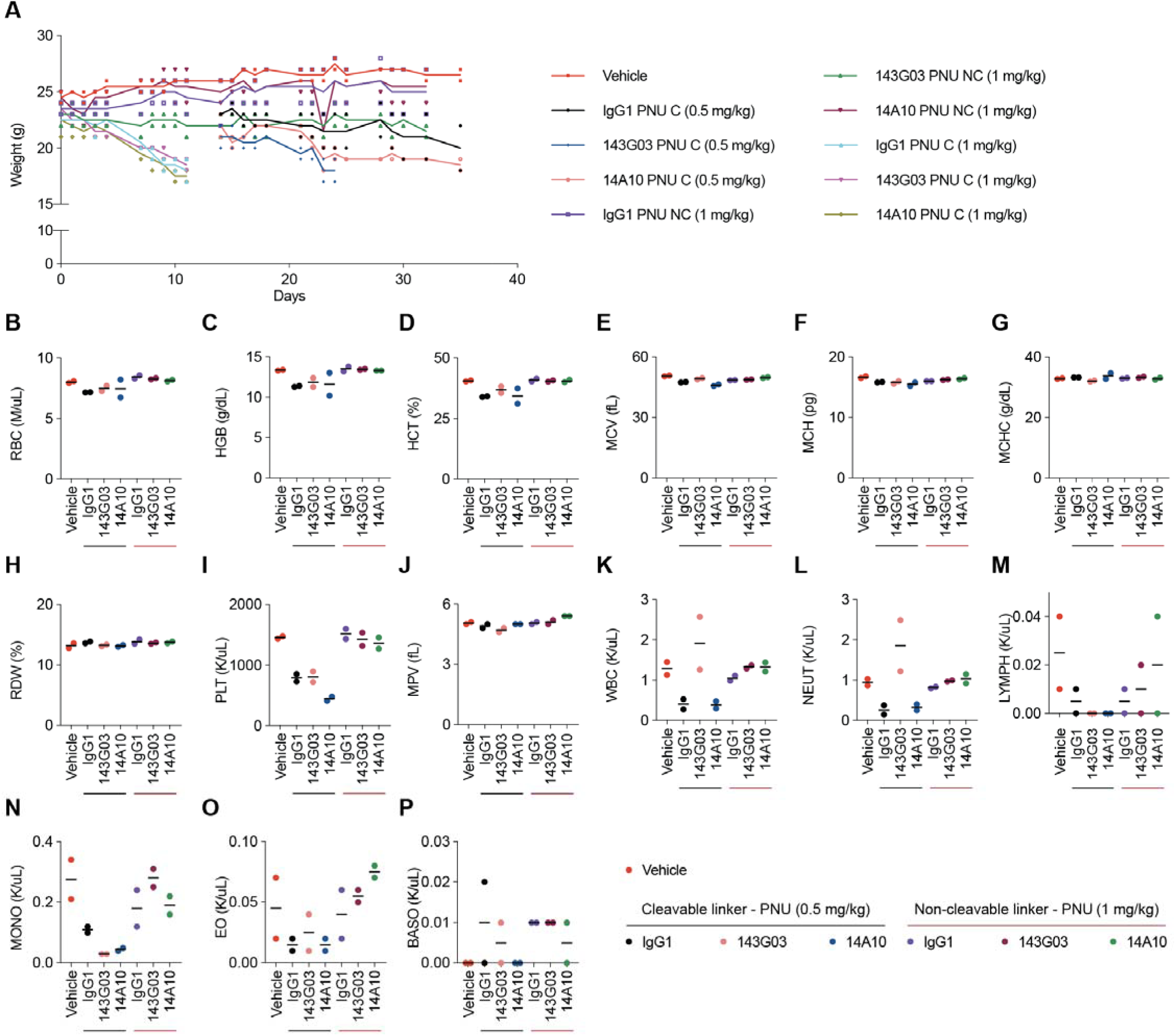
Safety assessment of *in vivo* L1CAM ADC treatments. **A**, NSG mice were administered weekly doses of the designated ADCs or vehicle control for a total of four weeks. Body weight was measured and monitored throughout the study as an indicator of adverse effects. *n* = 2 mice per treatment condition. C, cleavable; NC, non-cleavable linker. **B**-**P**, Complete blood count analysis of mice treated with ADCs or vehicle control at the time of sacrifice. Analyses show red blood cells, RBC (**B**), hemoglobin, HGB (**C**), hematocrit, HCT (**D**), mean corpuscular volume, MCV (**E**), mean corpuscular hemoglobin, MCH (**F**), mean corpuscular hemoglobin concentration, MCHC (**G**), red cell distribution width, RDW (**H**), platelet count, PLT (**I**), mean platelet volume, MPV (**J**), white blood cell count, WBC (K), neutrophil count, NEUT (**L**), lymphocyte count, LYMPH (**M**), monocyte count, MONO (**N**), eosinophil count, EO (**O**), and basophil count, BASO **(P**). *n* = 2 mice per treatment condition. Sample legends are shown in the bottom right corner.

**Supplementary Figure S2.**
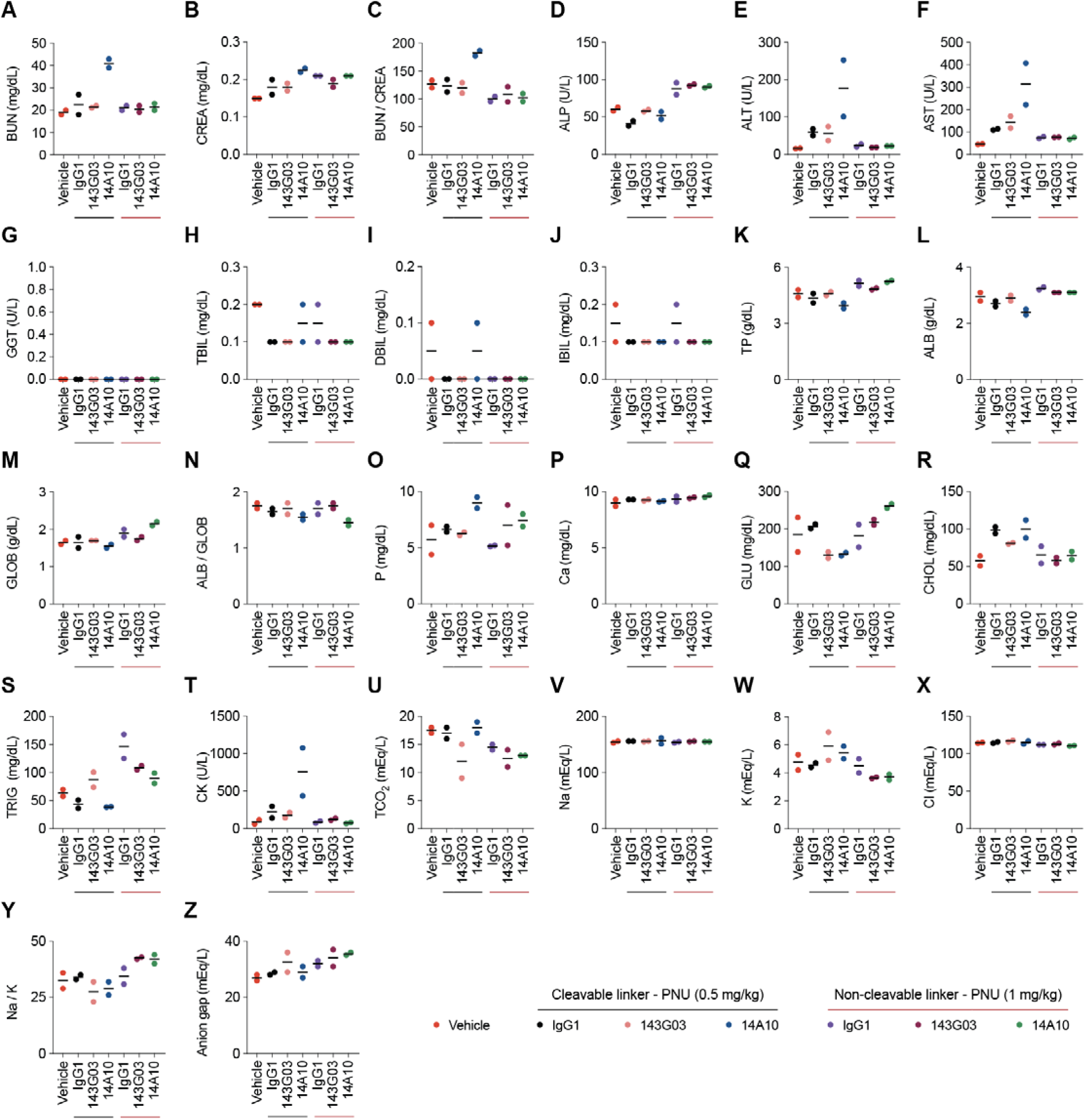
*In vivo* clinical chemistry assessment of mice treated with L1CAM ADCs. **A**-**Z**, Clinical chemistry analyses at the time of sacrifice. Analyses include blood urea nitrogen, BUN (**A**), creatinine, CREA (**B**), blood urea nitrogen to creatinine ratio (**C**), alkaline phosphatase, ALP (**D**), Alanine transferase, ALT (**E**), aspartate transferase, AST (**F**), gamma-glutamyl transferase, GGT (**G**), total bilirubin, TBIL (**H**), direct bilirubin, DBIL (**I**), indirect bilirubin IBIL (**J**), total protein, TP (**K**), albumin, ALB (**L**), globulin, GLOB (**M**), albumin to globulin ratio (**N**), phosphorus, P (**O**), calcium, Ca (**P**), glucose, GLU (**Q**), cholesterol, CHOL (**R**), triglycerides, TRIG (**S**), creatine kinase, CK (**T**), total carbon dioxide, TCO_2_ (**U**), sodium, Na (**V**), potassium, K (**W**), chlorine, Cl (**X**), sodium to potassium ratio (**Y**), and anion gap (**Z**). *n* = 2 mice per treatment condition. Sample legends are shown in the bottom right corner.

**Supplementary Figure S3.**
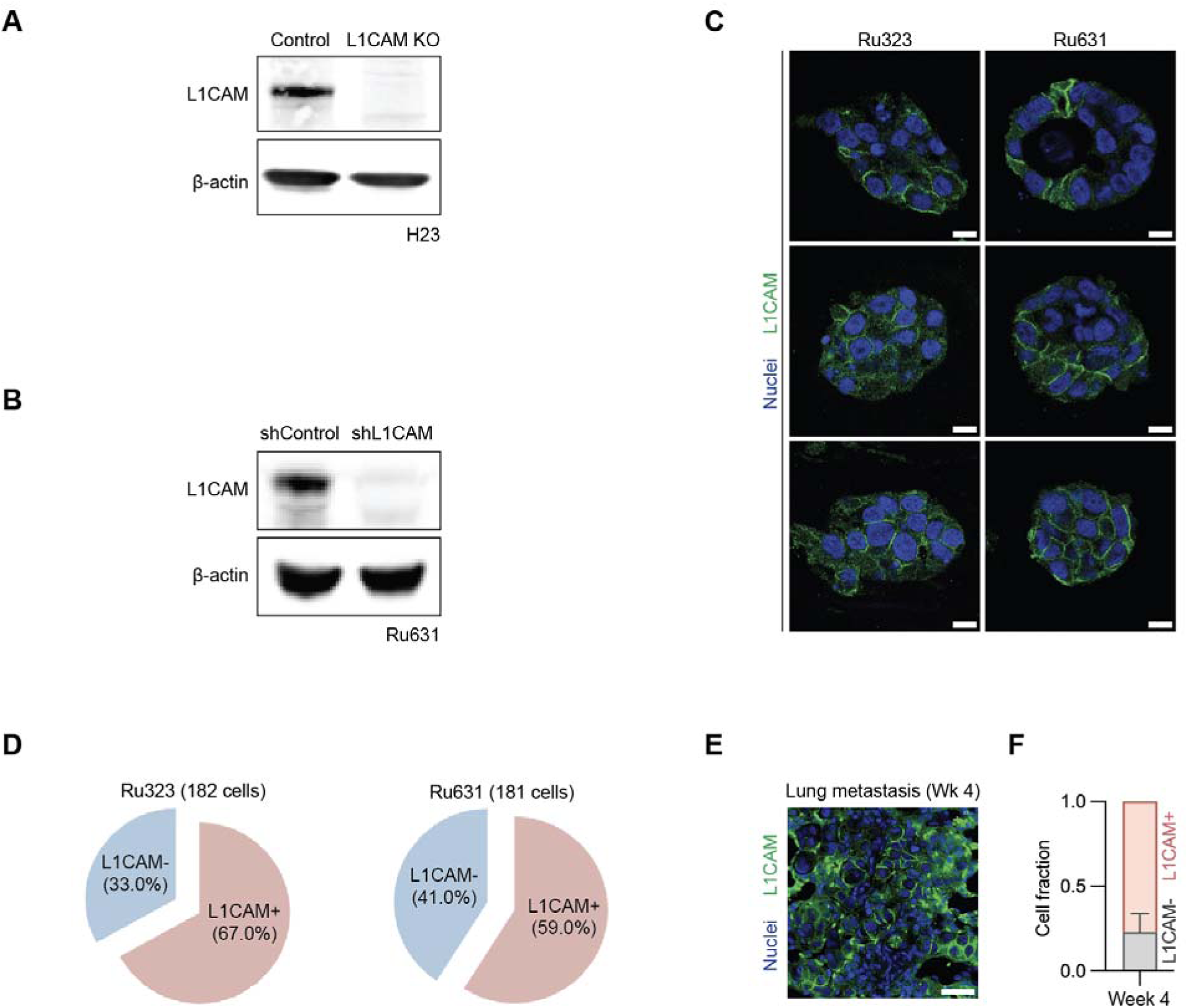
*In vivo* efficacy of L1CAM ADC against LUAD metastasis. **A**, L1CAM western immunoblotting anlysis in parental and L1CAM knockout H23 cells. **B**, Western immunoblotting analysis of L1CAM levels in Ru631 PDX tumoroids upon L1CAM knockdown. **C**, L1CAM IF with a nuclear counterstaining in Ru323 and Ru631 PDX tumoroids. Scale bar, 10 μm. **D**, Quantification of L1CAM^+^ cells in Ru323 PDX tumoroid (*n =* 182 cells) and Ru631 PDX tumoroid (*n =* 181 cells). **E**, L1CAM IF staining of lung tissue section with metastatic lesions generated four weeks after inoculation of Ru631 PDX cells into NSG mice via the tail vein. Scale bar, 40 μm. **F**, Quantification of L1CAM^−^ or L1CAM^+^ fraction of Ru631 PDX cells in the experiment of panel (**E**). *n* = 3 mice. Bar graph, mean ± S.D.

