## Supplementary table 1 for "Development of antibody-drug conjugates targeting L1CAM to treat metastatic cancer"

| mAb | Mouse | Epitope Bin | Epitope Domain | huL1CAM Protein $K_D$ [nM] | cyL1CAM Protein $K_D$ [nM] | huL1CAM 293T cells $EC_{50}$ [nM] | moL1CAM 293FT cells $EC_{50}$ [nM] | MCF-7 Cells $EC_{50}$ [nM] | MDA-231 Cells $EC_{50}$ [nM] |
|---|---|---|---|---|---|---|---|---|---|
| **13F04** | WT | A | Ig | 1.13 | 19.8 | 4 | NB | 1.2 | 1.7 |
| **143G03** | WT | C | FN | 0.12 | 0.34 | 1.0 | NB | 0.35 | 0.9 |
| **04B06** | AMM | D | FN | 0.07 | 0.6 | 2.3 | NB | 1.1 | 9.6 |
| **12D10** | AMM | C | FN | 0.15 | 1.9 | 1.2 | NB | 0.77 | 2 |
| **13G04** | AMM | D | FN | <0.02* | 3.2 | 4.8 | NB | 4.8 | 6.7 |
| **14A10** | AMM | B | Ig | 1.2 | 3.2 | 3.5 | 5.6 | 5.3 | 8.9 |
| **15G02** | AMM | E | FN | 0.58 | 0.93 | 2.6 | NB | 1.0 | 0.98 |

WT: wild-type mice; AMM: AlivaMab® Mice

Ig: immunoglobulin domain; FN: fibronectin domain

NB: no binding

*off-rate below detection limit

**Supplementary Table 1: Binding profile of candidate L1CAM mAbs**
